# Metal-organic framework (MOF) nanomedicine preparations of sildenafil designed for the future treatment of pulmonary arterial hypertension

**DOI:** 10.1101/718478

**Authors:** Nura A. Mohamed, Haissam Abou Saleh, Yu Kameno, Isra Marei, Gilberto de Nucci, Blerina Ahmetaj-Shala, Fisnik Shala, Nicholas S. Kirkby, Lewis Jennings, Robert P. Davies, Paul D. Lickiss, Jane A. Mitchell

**Author notes:** Authors for correspondence: Dr. Haissam Abou Saleh,: Dr. Nura Mohamed or Professor Jane A. Mitchell Department of Cardiothoracic Pharmacology, National Heart and Lung Institute, Imperial College, London, UK. Footnotes: sil@nanoMIL-89; sildenafil loaded nanoMIL-89.

## Abstract

Pulmonary Arterial Hypertension (PAH) is an aggressive disease with poor prognosis, no available cure, and low survival rates. Currently, there are several classes of vasodilator drugs that are widely used as treatment strategies for PAH. These include (i) endothelin-1 receptor antagonists, (ii) phosphodiesterase type 5 inhibitors, (iii) prostacyclin analogues, and (iv) soluble guanylate cyclase activators. Despite their clinical benefits, these therapies are hindered by their systemic side effects. This limitation could be overcome by controlled drug release, with future modifications for targeted drug delivery, using a nanomedicine approach. In the current study, we have evaluated one particular nanomedicine platform (the highly porous iron-based metal-organic framework (MOF) commonly referred to as MIL-89) as a carrier of the PAH drug sildenafil. We have previously shown that MIL-89 is relatively non-toxic in a range of human cell types and well tolerated *in vivo*. Here we prepared a nano-formulation of MIL-89 (nanoMIL-89) and then successfully charged it with a payload of sildenafil (sil@nanoMIL-89^1^). sil@nanoMIL-89 was then shown to release sildenafil in a biphasic manner with an initial rapid release over 6 hours followed by a more sustained release over 72 hours. Both nanoMIL-89 and sil@nanoMIL-89 were relatively non-toxic when incubated with human endothelial cells for 24 hours. Finally, the vasodilator effect of sil@nanoMIL-89 was measured over 8 hours using isolated mouse aorta. Consistent with drug release kinetics, sil@nanoMIL-89 induced vasodilation after a lag phase of >4 hours. Thus, in sil@nanoMIL-89, we have produced a nano-formulation of sildenafil displaying delayed drug release corresponding to vasodilator activity. Further pharmacological assessment of sil@nanoMIL-89, including in PAH models, is now required and constitutes the subject of ongoing investigation.

## Introduction

Pulmonary arterial hypertension (PAH) is a devastating disease within which pulmonary arteries constrict and remodel, resulting in elevated pulmonary artery pressure and increased workload on the right side of the heart. This ultimately leads to right heart failure and premature death. Whilst there is no cure for PAH, there are four classes of vasodilator drugs currently used to help slow disease progression. These drugs include; (i) phosphodiesterase type 5 inhibitors, such as sildenafil, (ii) soluble guanylate cyclase activator drugs, such as riociguat, (iii) endothelin-1 receptor antagonists, such as bosentan and (iv) prostacyclin analogs such as treprostinil sodium. Whilst these drugs show efficacy in PAH, they have limited pharmacokinetics and affect the systemic circulation, which ultimately limits the dose of drug that can be used. As such, we[1-3] and others[4-6] have suggested that PAH is a disease that would benefit from the application of controlled drug release, with the possibility of introducing targeted drug delivery strategies, using a nanomedicine approach. Whilst this idea is relatively novel, a limited number of studies are emerging describing nanomedicine formulations suitable for application in PAH. These include (i) a liposome-conjugate that combines the Rho-kinase inhibitor and vasodilator drug fasudil with the nitric oxide (NO) donor DETA NONOate[7], (ii) a liposome-encapsulated Iloprost nano-formulation[8] and from our own group, a polymeric NO-releasing nanoparticle[3]. Unfortunately liposomes and polymers as nano-formulation platforms have limitations including: (i) difficulty of *in vivo* imaging without further chemical modification, (ii) low solubility window, (iii) difficulty of drug fusion and encapsulation, (iv) high production cost and (v) difficulty in maintaining stability and bioactivity of drugs during the conjugation process[9,10]. We have previously highlighted the potential of nanoscale metal-organic frameworks (MOFs; nanoMOFs) as carriers for PAH drugs. In this regard, MOFs have several advantages as drug delivery platforms including (i) being biocompatible and biodegradable, (ii) high thermal, chemical and mechanical stability, (iii) control over nanoparticle size, (iv) extremely high porosities with commensurate high drug loading capacity and (v) the ability to tailor the size, shape and chemical nature of the internal and external surfaces thus allowing a high degree of control over drug-binding and release kinetics[11,12]. We have focussed our initial work on the iron-containing MOF, MIL-89 since this particular material has the added advantage of potentially being imageable using magnetic resonance imaging (MRI)[13,14] and a predicted cavity/pore size that is suitable for encapsulating PAH drugs[2]. In our hands, MIL-89 was relatively non-toxic in a range of human cell types, including endothelial and vascular smooth muscle cells and was well tolerated *in vivo* for two weeks[2]. In line with this, others have found that a related iron-based MOF was well tolerated *in vivo* for up to three months[15].

Here we have extended our studies by firstly preparing sildenafil-loaded nanoMIL-89 (sil@nanoMIL-89^1^) and then studying its drug release performance and corresponding vasodilator function.

## Material and Methods

### Preparation of nanoMIL-89

NanoMIL-89 was prepared following a modification of the previously published procedure by Horcajada et al.,[16] with the addition of an optimised quantity of glacial acetic acid to the reaction mixture to control particle size and quality. In brief, 10 mmol of iron(III) chloride hexahydrate (FeCl3.6H2O; Sigma Aldrich, UK) and 10 mmol *trans, trans*-muconic acid (Sigma Aldrich, UK) were mixed in absolute ethanol (100 ml; 99.8%; VWR, UK). The reaction mixture was then sonicated for 15 minutes, and 20 ml of glacial acetic acid (99.8%; VWR, UK) was added. The mixture was heated at 90°C for 24 hours in a Parr reactor after which the precipitate was recovered by centrifugation at 7000 rpm for 15 min. The precipitate was washed with distilled water and vacuum dried to recover the nanoMIL-89 which presented as a brown precipitate (100-200 mg/reaction).

### Chemical characterization of nanoMIL-89

The successful preparation of nanoMIL-89 was confirmed using powder X-ray diffraction (PXRD; Bruker D2 Phaser) studies and infrared/attenuated total reflection (IR/ATR; Perkin Elmer Spectrum). Particle size was measured using dynamic light scattering (DLS; DelsaNano C by Beckman Coulter) where 50 μl of the reaction mixture was taken and added to 5 ml of ethanol which was then centrifuged for 2 minutes before 100 μl of the supernatant was used for DLS analysis. In addition, particle size was estimated from images obtained from dried nanoMIL-89 samples using scanning electron microscopy (SEM; LEO Gemini 1525 by Zeiss), analysed and quantified using Image J Launcher Software[17] to identify particulate size. Briefly, 20 nanoparticles were selected randomly with the only prerequisite being that they be well defined. Length and width measurements were made using ImageJ. PXRD analysis was consistent with literature patterns for bulk MIL-89 (See Figure 1).[2,18,19]

**Figure 1:**
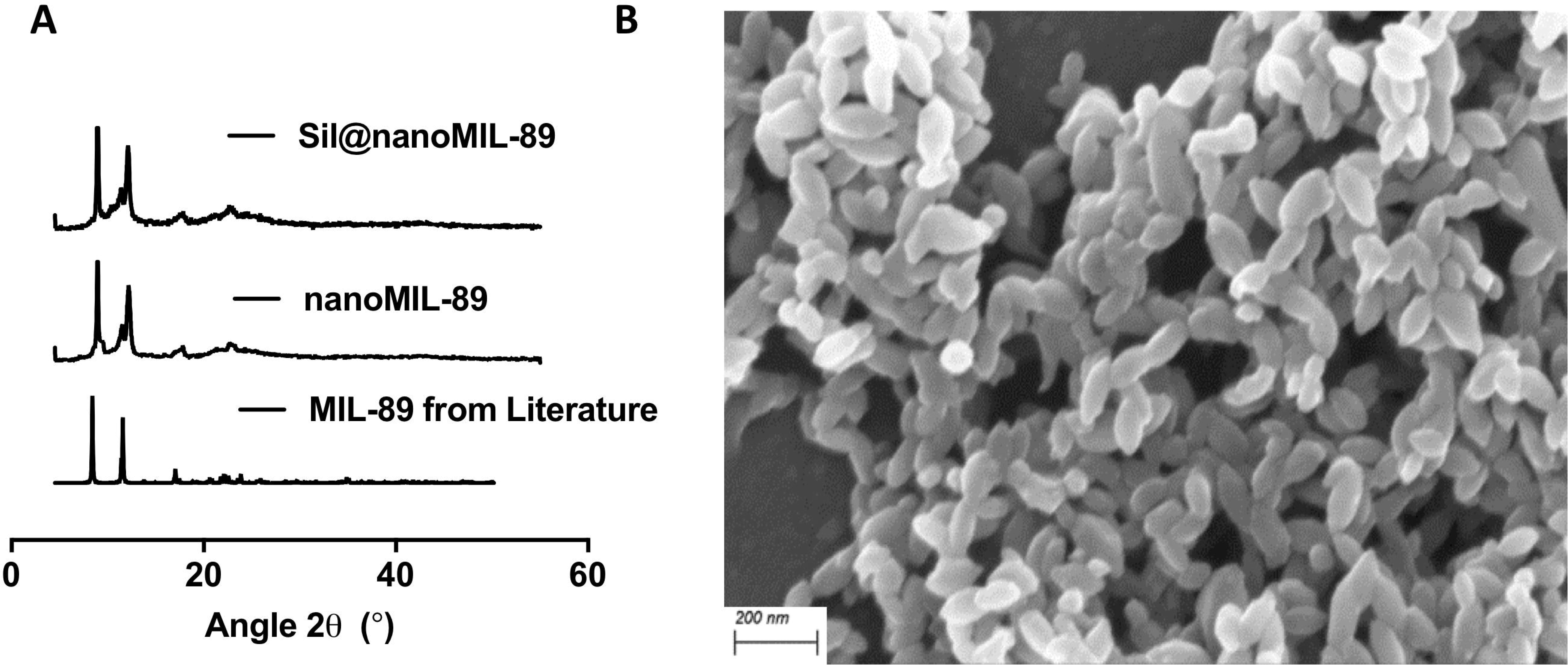
Characterization of nanoMIL-89. (A) Powder X-ray Diffraction (PXRD) analysis of nanoMIL-89 prepared within the reported study, MIL-89 reported in literature[12] and nanoMIL-89 loaded with sildenafil (sil@nanoMIL-89). (B) Scanning Electron Microscope image of nanoMIL-89 (x5,000).

### Loading and release studies of nanoMIL-89 with sildenafil

The sildenafil nanoMIL-89 loading procedure, to produce sil@nanoMIL-89, was conducted as follows: 5 ml of a 1 mg/ml sildenafil solution was prepared in PBS (pH 7.4; Sigma Aldrich, UK), then 20 mg of nanoMIL-89 was added to the solution and incubated on a shaker at room temperature for 16-18 hours. The concentration of sildenafil was selected to be just below its maximum solubility in aqueous solutions. The resultant sil@nanoMIL-89 precipitate was retrieved by centrifugation at 7000 rpm at room temperature for 15 minutes. Samples of the supernatant were collected to assess levels of sildenafil remaining after loading and were used to calculate the amount of drug taken up by the nanoMIL-89. The precipitated sil@nanoMIL-89 was then re-suspended in 5 ml of human plasma from 3 separate donors (Cambridge Bioscience, UK). Solutions of sil@nanoMIL-89 were incubated at 37°C for 0, 0.5, 1, 3, 6, 16, 24, 36, 48, 60, 72, 84 and 96 hours. At these time points, the solution was centrifuged at 7000 rpm at room temperature for 15 minutes. The release of sildenafil into the supernatant was measured at each time point by ELISA (MaxSignal® Sildenafil/Vardenafil ELISA Test Kit (MEDIBENA, UK) following the manufacturer’s instructions.

### Human endothelial cell viability response to sil@nanoMIL-89

Human blood outgrowth endothelial cells (BOECs) were isolated, cultured, plated, and treated as described previously[2]. Cell viability was indicated by changes in respiration using alamarBlue™ reagent according to manufactures instructions.

### Aorta vasomotor responses to sil@nanoMIL-89

C57 Black 4 mice (6-10 weeks) were killed by CO_2_ narcosis and aorta removed, cleaned of connective tissue and cut into 1.5 mm rings before being mounted in Mulvany-Halpern myograph organ baths containing a physiological salt solution (PSS), as we have described previously[20]. In order to optimize the stability of vascular function over the 8-hour time course, diclofenac (1 *µ*M) and cycloheximide (1 *µ*M) were added to the PSS to block vasoactive prostanoids and induction of vasoactive genes (e.g. NO synthase) respectively. Vessels were contracted with an EC_80_ concentration of U46619 (10 nM). Once a stable baseline was obtained, sildenafil (10 *µ*M), nanoMIL-89 (10 *µ*g/ml) or sil@nanoMIL-89 (10 *µ*g/ml) was added to the PSS, and vascular tone monitored for 8 hours. Responses in vessels incubated in PSS served as controls. Force was recorded via a PowerLab/800 (AD Instruments Ltd., UK) and analysed using Chart 6.0 acquisition system (AD Instruments Ltd.,UK).

### Statistical analysis

Data were presented as mean ± SEM and statistical significance (taken as P < 0.05) was determined using GraphPad Prism 7 as described in each figure legend.

## Results and Discussion

### Chemical characterization of nanoMIL-89

NanoMIL-89 was prepared following previously reported procedures for MIL-89[2,16] but with an additional step of the inclusion of an optimised quantity of glacial acetic acid to the reaction mixture. Adjusting the amount of glacial acetic acid allowed control over the nanoparticle size and size distribution. The successful preparation of nanoMIL-89 was confirmed using PXRD analysis (Figure 1A) and IR/ATR (Supplementary Figure 1). Particle size was measured using DLS (Supplementary Figure 2) and SEM (Figure 1B). SEM analysis of solid nanoMIL-89 revealed ovoid nanoparticles of length 82.5±20.2 nm and width 31.4±6.6 nm (n=20) consistent with the 50-100 nm particle size reported in the original study by Horcajada et al[16]. DLS analysis of 3 consecutive batches of nanoMIL-89 in solvent showed an estimated gyration diameter of 87.6±25.3 nm (Supplementary Figure 2), 137.1±40.3 nm and 147.6±17.98 nm. These results confirm that nanoMIL-89 prepared in the current study was in the appropriate nanoscale range for biological applications (i.e. <150 nm) and conforms to the expected composition and structure. The aim of this study was to extend our previous work[2] and load nanoMIL-89 with a relevant PAH drug, namely sildenafil. The stability of the nanoMOF was confirmed by PXRD studies after suspending it overnight (16-18 hours) in medium both in the absence and presence of sildenafil. Moreover, the PXRD pattern of nanoMIL-89 and sil@nanoMIL-89 were similar, indicating that the structure retained its integrity on loading (Figure 1A).

### Sildenafil loading and release studies

Loading studies showed that, under our experimental conditions, nanoMIL-89 readily absorbed >90% of the sildenafil in the loading solution to yield sil@nanoMIL-89. This on a mass basis amounted to 17% of the starting weight of nanoMIL-89. The high efficiency of sildenafil uptake from the loading solution and high loading wt% are both promising for potential medical applications. In addition, since the uptake of sildenafil from the loading solution was virtually complete, it is possible that even higher loading capacities can be achieved. Sildenafil release from sil@nanoMIL-89 was then measured using human plasma as a matrix. As expected, plasma incubated with nanoMIL-89 did not react with reagents in the sildenafil ELISA (Figure 2). Authentic sildenafil was relatively stable in plasma for up to 48 hours, after which it completely degraded to undetectable levels. sil@nanoMIL-89 exhibited an early steady release of sildenafil with detectable levels present in plasma within the first hour. After 6 hours, sil@nanoMIL-89 was still actively releasing the drug, although at a reduced rate (Figure 2). Interestingly, at the 72 and 96 hour time points when free sildenafil(i.e. in the absence of the MOF) had degraded, the sil@nanoMIL-89 sample continued to show high and consistent levels of sildenafil due to its continuing slow release (Figure 2). The maximum amount of sildenafil released by sil@nanoMIL-89 in plasma corresponds to 51% of the originally loaded drug, although the actual value could be higher due to degradation of the sildenafil in plasma over time.

**Figure 2:**
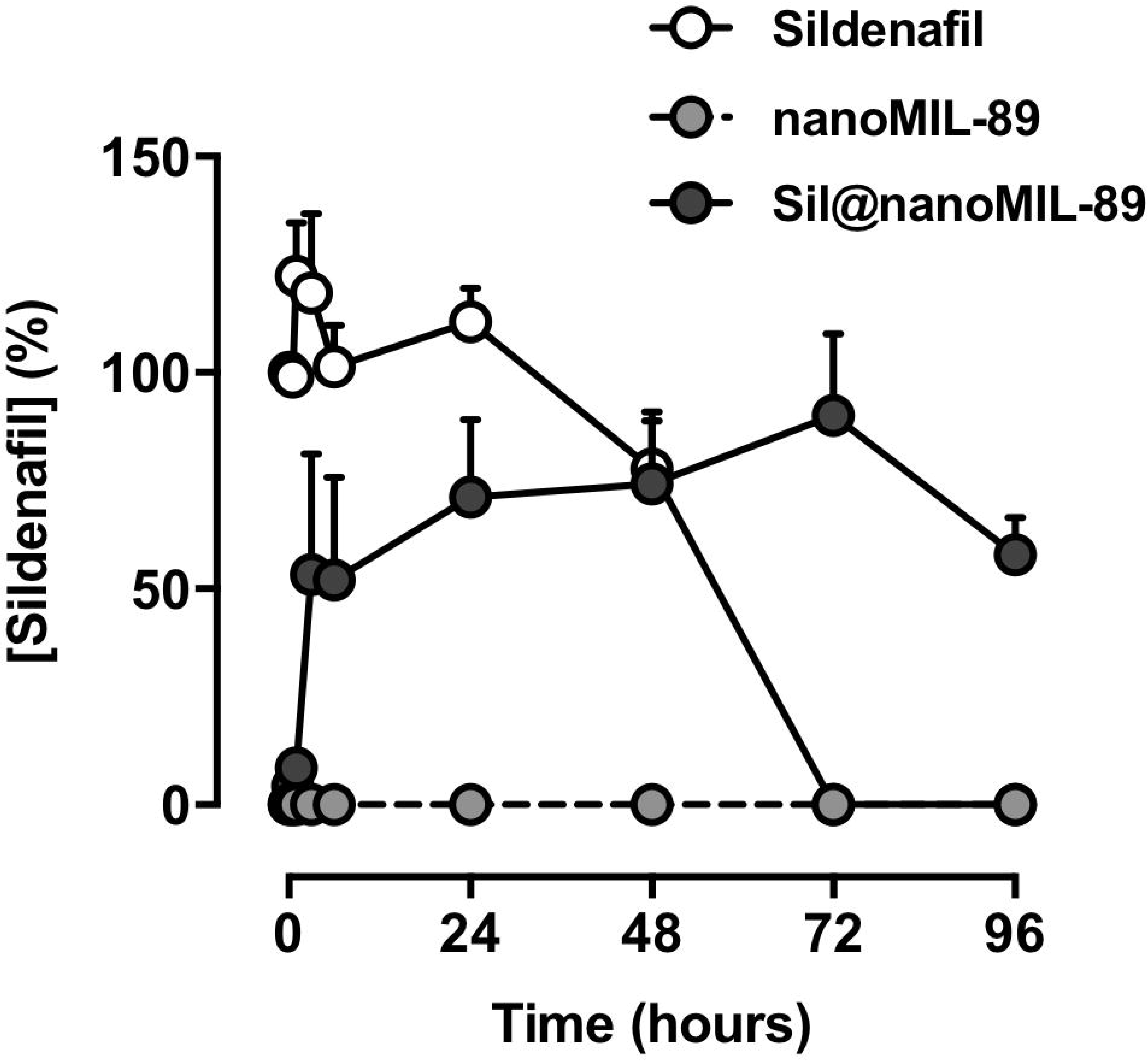
Sildenafil release by sil@nanoMIL-89. Sildenafil levels in plasma were measured from incubations of sil@nanoMIL-89 (4 mg/ml), nanoMIL-89 (4 mg/ml) and sildenafil (1 mg/ml) at 37°Cover 96 hours. Data are mean ± SEM for n=6.

### Effect of nanoMIL-89 and sil@nanoMIL-89 on cell viability

As we have shown previously for nanoMIL-89[2] and in the current study for sil@nanoMIL-89, the viability of human BOECs was not significantly affected by either MOF incubated with cells for 24 hours (Figure 3).

**Figure 3:**
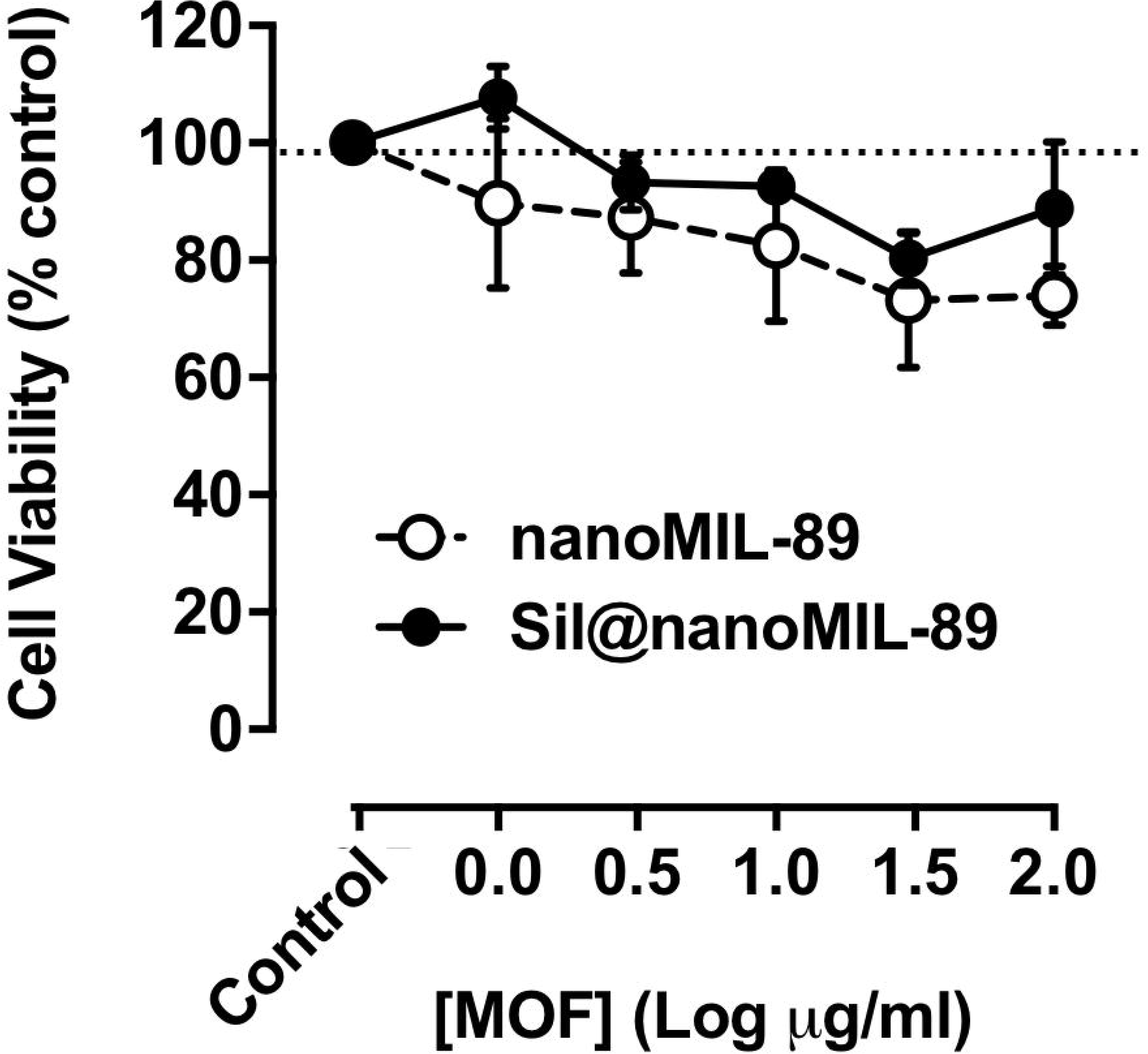
Effect of nanoMIL-89 and sil@nanoMIL-89 on cell viability in human blood outgrowth endothelial cells. Data are shown as mean ± SEM for n=8 determinations using cells from 4 separate isolations.

### Vasodilator responses of Sil-nanoMIL-89

Over the course of 8 hours mouse aorta contracted with U46619, the synthetic mimetic of the endoperoxide prostaglandin PGH_2_ which acts on thromboxane receptors in smooth muscle cells, retained approximately 70% of induced tone (Figure 4). Sildenafil is a vasodilator drug which relaxes blood vessels by increasing the biological half-life of the cGMP. As expected, therefore, sildenafil-induced vasodilation within the first 60 minutes and continued to relax vessels (compared to control) for the duration of the experiment (Figure 4). NanoMIL-89 also induced relaxation, but with a different time course, relaxation induced by nanoMIL-89 was not apparent (compared to PSS alone) until >4 hours after addition of the MOF (Figure 4). It is not clear why nanoMIL-89 induced relaxation, but one explanation might be that the MOF absorbed the contractile agent U46619. sil@nanoMIL-89 induced clear and sustained vasodilator responses, which began after a lag phase of >4 hours and were sustained for the duration of the experiment (Figure 4). The maximal relaxant effect of sil@nanoMIL-89 (10 *µ*g/ml) was greater than either nanoMIL-89 (10 *µ*g/ml) or sildenafil alone (10 *µ*M) (Figure 4).

**Figure 4:**
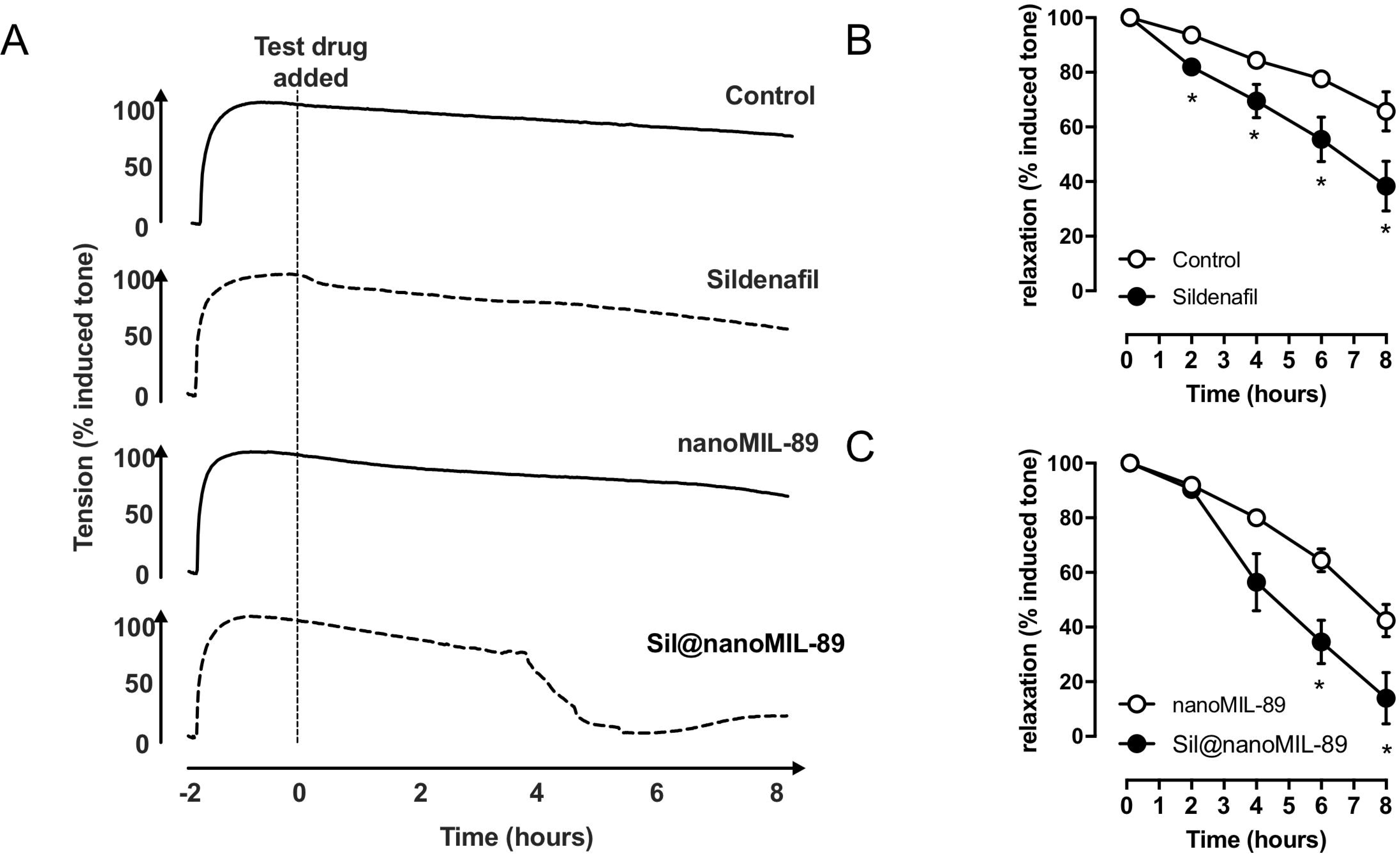
Vasodilator effects of sildenafil (10 *µ*M), nanoMIL-89 (10 *µ*g/ml), and sil@nanoMIL-89 (10 *µ*g/ml) on pre-contracted mouse aorta. The figure shows representative vessel responses across the full time course (A) and pooled data presented as mean ± SEM for n=3-8 vessels from 3-4 mice (B and C). Statistical significance was determined by two-way ANOVA followed by Tukey’s multiple comparisons test. Statistical significance was assumed where *p<0.05.

## Conclusion

Here we have prepared, for the first time, a sildenafil-loaded nanoMOF material and assessed its potential in drug delivery applications. sil@nanoMIL-89 retained its crystallinity, released sildenafil over a prolonged period of time, and induced vasodilation in a sustained manner. These results are consistent with the idea that nanoMIL-89 is a promising formulation prototype for PAH drugs and describe our first prototype drug (sil@nanoMIL-89). The next steps will be (i) to assess the pharmacology, toxicology, efficacy and *in vivo* distribution of sil@nanoMIL-89 and (ii) establish strategies for targeted delivery of sil@nanoMIL-89 and related drugs to the pulmonary vasculature.

## Acknowledgements

The authors are indebted to Ms Hime Gashaw for her expert training in cell culture techniques.

## Figure legends

**Supplementary Figure 1: Infrared/attenuated total reflection (IR/ATR) spectra for nanoMIL-89**. The figure shows a representative IR/ATR tracing.

**Supplementary Figure 2: Estimation of nanoMIL-89 size using dynamic light scattering (DLS) analysis**.

## References

[1] J.A. Mitchell, B. Ahmetaj-Shala, N.S. Kirkby, W.R. Wright, L.S. Mackenzie, D.M. Reed, N. Mohamed, Role of prostacyclin in pulmonary hypertension, Glob Cardiol Sci Pract 2014 (2014) 382-393. 10.5339/gcsp.2014.53.

[2] N.A. Mohamed, R.P. Davies, P.D. Lickiss, B. Ahmetaj-Shala, D.M. Reed, H.H. Gashaw, H. Saleem, G.R. Freeman, P.M. George, S.J. Wort, D. Morales-Cano, B. Barreira, T.D. Tetley, A.H. Chester, M.H. Yacoub, N.S. Kirkby, L. Moreno, J.A. Mitchell, Chemical and biological assessment of metal organic frameworks (MOFs) in pulmonary cells and in an acute in vivo model: relevance to pulmonary arterial hypertension therapy, Pulm Circ 7 (2017) 643-653. 10.1177/2045893217710224.

[3] N.A. Mohamed, B. Ahmetaj-Shala, L. Duluc, L.S. Mackenzie, N.S. Kirkby, D.M. Reed, P.D. Lickiss, R.P. Davies, G.R. Freeman, B. Wojciak-Stothard, A.H. Chester, I.M. El- Sherbiny, J.A. Mitchell, M.H. Yacoub, A New NO-Releasing Nanoformulation for the Treatment of Pulmonary Arterial Hypertension, J Cardiovasc Transl Res 9 (2016) 162-164. 10.1007/s12265-016-9684-2.

[4] V. Segura-Ibarra, S. Wu, N. Hassan, J.A. Moran-Guerrero, M. Ferrari, A. Guha, H. Karmouty-Quintana, E. Blanco, Nanotherapeutics for Treatment of Pulmonary Arterial Hypertension, Front Physiol 9 (2018) 890. 10.3389/fphys.2018.00890.

[5] J.S. Brenner, C. Greineder, V. Shuvaev, V. Muzykantov, Endothelial nanomedicine for the treatment of pulmonary disease, Expert Opin Drug Deliv 12 (2015) 239-261. 10.1517/17425247.2015.961418.

[6] W. Mosgoeller, R. Prassl, A. Zimmer, Nanoparticle-mediated treatment of pulmonary arterial hypertension, Methods in enzymology 508 (2012) 325-354. 10.1016/b978-0-12-391860-4.00017-3.

[7] J. Rashid, K. Nahar, S. Raut, A. Keshavarz, F. Ahsan, Fasudil and DETA NONOate, Loaded in a Peptide-Modified Liposomal Carrier, Slow PAH Progression upon Pulmonary Delivery, Molecular pharmaceutics 15 (2018) 1755-1765. 10.1021/acs.molpharmaceut.7b01003.

[8] P.P. Jain, R. Leber, C. Nagaraj, G. Leitinger, B. Lehofer, H. Olschewski, A. Olschewski, R. Prassl, L.M. Marsh, Liposomal nanoparticles encapsulating iloprost exhibit enhanced vasodilation in pulmonary arteries, International journal of nanomedicine 9 (2014) 3249- 3261. 10.2147/ijn.s63190.

[9] A. Akbarzadeh, R. Rezaei-Sadabady, S. Davaran, S.W. Joo, N. Zarghami, Y. Hanifehpour, M. Samiei, M. Kouhi, K. Nejati-Koshki, Liposome: classification, preparation, and applications, Nanoscale research letters 8 (2013) 102. 10.1186/1556-276x-8-102.

[10] J.C. Erik Brewer, and Anthony Lowman, Emerging Technologies of Polymeric Nanoparticles in Cancer Drug Delivery, Journal of Nanomaterials Volume 2011 (2011) 10.

[11] S.K.a.S. Kızılel, Biomedical Applications of Metal Organic Frameworks, Industrial & Engineering Chemistry Research, 2011 50 (4), 1799–1812.

[12] P. Horcajada, C. Serre, M. Vallet-Regi, M. Sebban, F. Taulelle, G. Ferey, Metal-organic frameworks as efficient materials for drug delivery, Angewandte Chemie (International ed. in English) 45 (2006) 5974-5978. 10.1002/anie.200601878.

[13] J. Estelrich, M.J. Sanchez-Martin, M.A. Busquets, Nanoparticles in magnetic resonance imaging: from simple to dual contrast agents, International journal of nanomedicine 10 (2015) 1727-1741. 10.2147/ijn.s76501.

[14] M.R. Prince, H.L. Zhang, S.G. Chabra, P. Jacobs, Y. Wang, A pilot investigation of new superparamagnetic iron oxide (ferumoxytol) as a contrast agent for cardiovascular MRI, Journal of X-ray science and technology 11 (2003) 231–240.

[15] A.A. Simagina, Towards rational design of metal-organic framework-based drug delivery systems, in: M.V. Polynski (Ed.), Russian Academy of Sciences and Turpion Ltd, Russia, 2018.

[16] P. Horcajada, T. Chalati, C. Serre, B. Gillet, C. Sebrie, T. Baati, J.F. Eubank, D. Heurtaux, P. Clayette, C. Kreuz, J.S. Chang, Y.K. Hwang, V. Marsaud, P.N. Bories, L. Cynober, S. Gil, G. Ferey, P. Couvreur, R. Gref, Porous metal-organic-framework nanoscale carriers as a potential platform for drug delivery and imaging, Nature materials 9 (2010) 172-178. 10.1038/nmat2608.

[17] C.A. Schneider, W.S. Rasband, K.W. Eliceiri, NIH Image to ImageJ: 25 years of image analysis, Nature methods 9 (2012) 671–675.

[18] C. Serre, S. Surble, C. Mellot-Draznieks, Y. Filinchuk, G. Ferey, Evidence of flexibility in the nanoporous iron(iii) carboxylate MIL-89, Dalton Trans (2008) 5462-5464. 10.1039/b805408h.

[19] S. Surble, F. Millange, C. Serre, G. Ferey, R.I. Walton, An EXAFS study of the formation of a nanoporous metal-organic framework: evidence for the retention of secondary building units during synthesis, Chem Commun (Camb) (2006) 1518-1520. 10.1039/b600709k.

[20] B. Ahmetaj-Shala, N.S. Kirkby, R. Knowles, M. Al’Yamani, S. Mazi, Z. Wang, A.T. Tucker, L. Mackenzie, P.C. Armstrong, R.M. Nusing, J.A. Tomlinson, T.D. Warner, J. Leiper, J.A. Mitchell, Evidence that links loss of cyclooxygenase-2 with increased asymmetric dimethylarginine: novel explanation of cardiovascular side effects associated with anti-inflammatory drugs, Circulation 131 (2015) 633-642. 10.1161/CIRCULATIONAHA.114.011591.

