## Supplementary figures and images for "Metal-organic framework (MOF) nanomedicine preparations of sildenafil designed for the future treatment of pulmonary arterial hypertension"

### Supplementary Figure 1

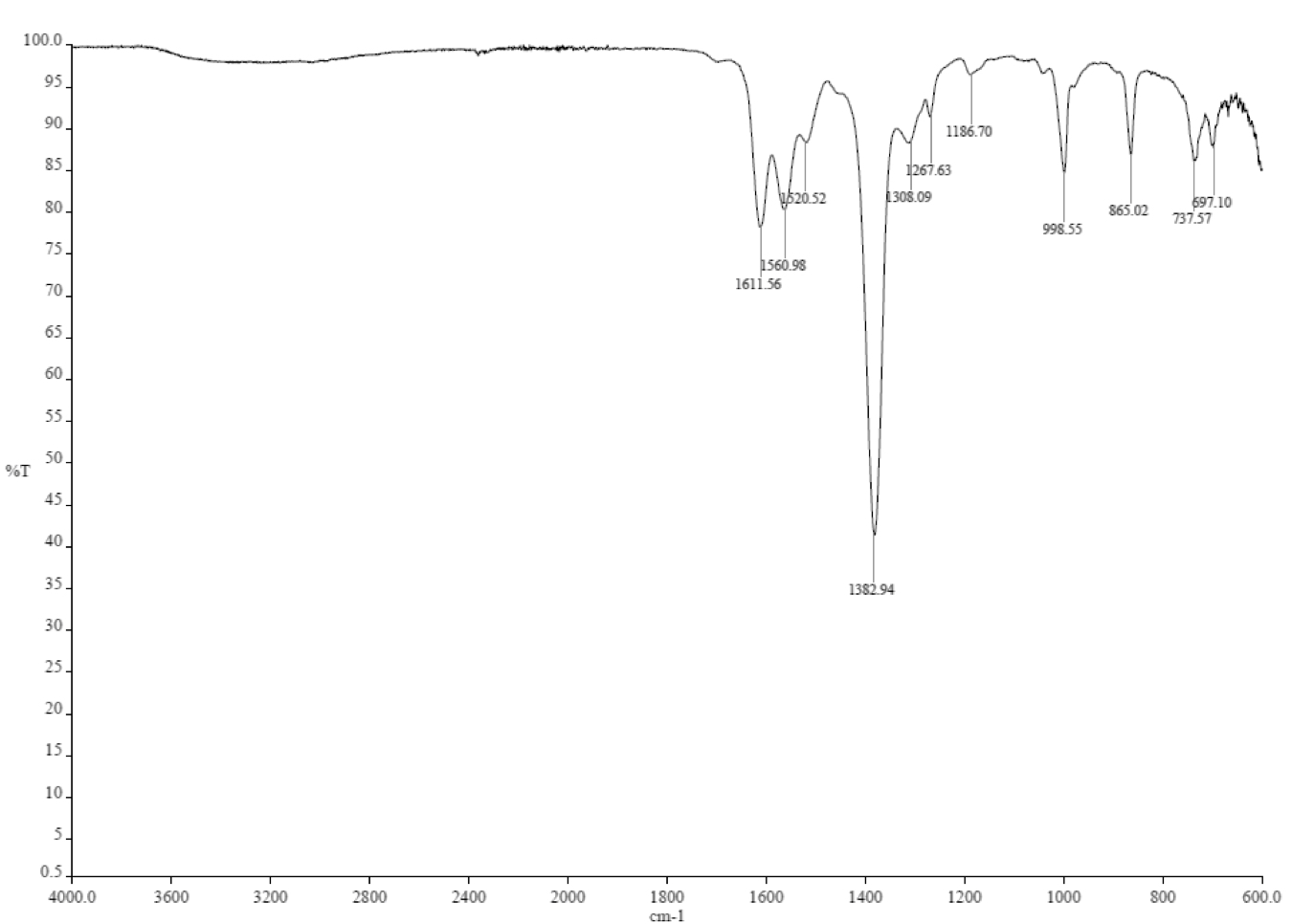

### Supplementary Figure 2

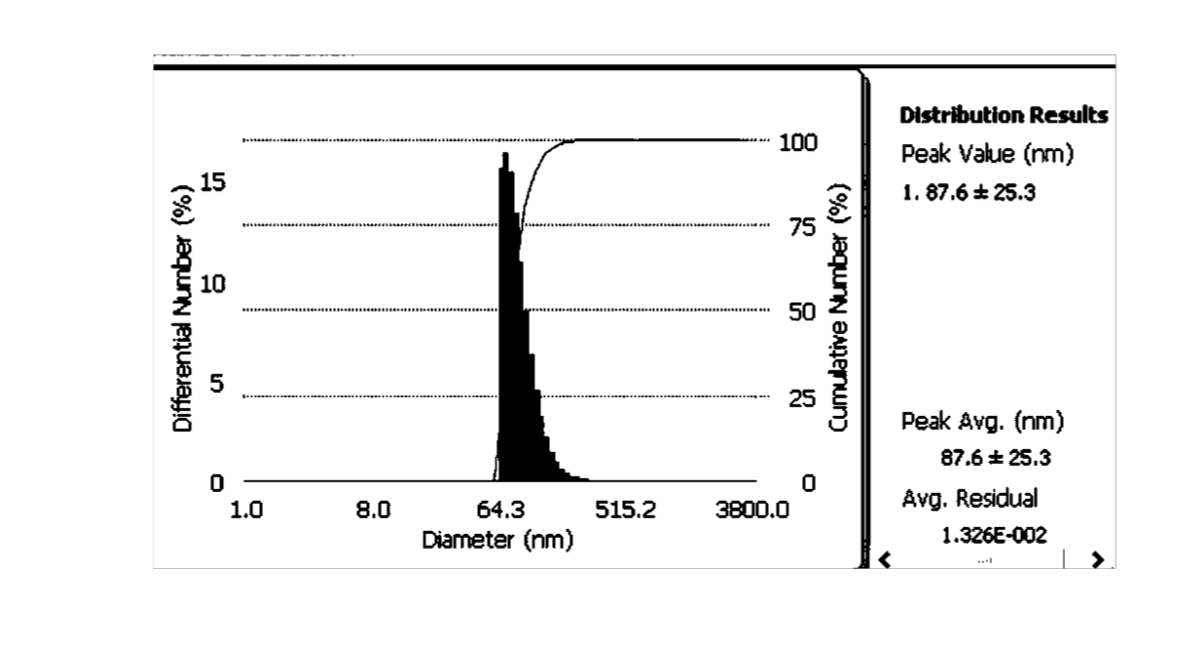
